# Efficacy of a Growth Hormone-Releasing Hormone Agonist in a Murine Model of Cardiometabolic Heart Failure with Preserved Ejection Fraction

**DOI:** 10.1101/2022.09.18.508429

**Authors:** Rosemeire M. Kanashiro-Takeuchi, Lauro M. Takeuchi, Raul A. Dulce, Katarzyna Kazmierczak, Wayne Balkan, Renzhi Cai, Wei Sha, Andrew V. Schally, Joshua M. Hare

## Abstract

Heart failure (HF) with preserved ejection fraction (HFpEF) represents a major unmet medical need owing to its diverse pathophysiology and lack of effective therapies. Potent synthetic, agonists (MR-356 and MR409) of growth hormone-releasing hormone (GHRH) improve the phenotype of models of HF with reduced ejection fraction (HFrEF) and in cardiorenal models of HFpEF. Endogenous GHRH exhibits a broad range of regulatory influences in the cardiovascular (CV) system, aging and plays a role in several cardiometabolic conditions including obesity and diabetes. Whether agonists of GHRH can improve the phenotype of cardiometabolic HFpEF remains untested and unknown. Here we tested the hypothesis that MR-356 can mitigate/reverse the cardiometabolic HFpEF phenotype. C57BL6N mice received a high fat diet (HFD) plus the nitric oxide synthase inhibitor (L-NAME) for 9 weeks. After 5 weeks of HFD+L-NAME regimen, animals were randomized to receive daily injections of MR-356 or placebo during a 4-week period. Control animals received no HFD+L-NAME or agonist treatment. Our results showed the unique potential of MR-356 to treat several HFpEF-like features including cardiac hypertrophy, fibrosis, capillary rarefaction, and pulmonary congestion. MR-356 improved cardiac performance by improving diastolic function, global longitudinal strain (GLS), and exercise capacity. Importantly, the increased expression of cardiac pro-brain natriuretic peptide (pro-BNP), inducible nitric oxide synthase (iNOS) and vascular endothelial growth factor-A (VEGF-A) was restored to normal levels suggesting that MR-356 reduced myocardial stress associated with metabolic inflammation in HFpEF. Thus, agonists of GHRH may be an effective therapeutic strategy for the treatment of cardiometabolic HFpEF phenotype.

## Introduction

Heart failure (HF) with preserved ejection fraction (HFpEF) is increasing in incidence and is becoming the leading form of HF worldwide (1, 2). HFpEF is associated with major cardiovascular (CV) risk factors such as aging, hypertension, obesity, and diabetes. To address the clinical burden imposed by HFpEF, new mechanistic insights and treatment options are urgently needed, but are very limited, due in large part to the complex pathogenesis involving multiple organs (lung, kidney, and skeletal muscle). The paucity of preclinical disease models that fully recapitulate the diversity of HFpEF phenotypes (3-5) is an additional challenge to conducting mechanistic studies that elucidate pathophysiology and provide insights for designing novel and effective therapeutic approaches tailored to improve cardiac performance in patients with HFpEF.

We previously showed that agonists of GHRH of the MR series (MR-356, MR-409) are cardioprotective in murine (6, 7) and porcine (8) models of HF with reduced ejection fraction (HFrEF) and other cardiovascular conditions such as vascular calcification (9) and non-ischemic cardiomyopathy (10). Recently, we demonstrated that GHRH-agonists have beneficial effects in cardiorenal HFpEF models: the murine HFpEF phenotype induced by chronic administration of angiotensin-II (Ang-II) (11) and in a porcine chronic kidney disease (CKD)-induced HFpEF model (12).

GHRH agonists exhibit a broad range of regulatory influences in the CV system, independent of their role in the GH-IGF-I axis (10, 13, 14), and in several cardiometabolic conditions including obesity (15) and diabetes (16). Administration of a GHRH-agonist to obese patients produced significant benefits, including reduced visceral adiposity, triglycerides, and measures of cardiovascular risk, with no changes in insulin sensitivity (15). Preclinical and clinical studies demonstrate the potential therapeutic benefits of GHRH-agonists for diabetes (17-20). Therefore, we focused our investigation on determining whether MR-356 can specifically target the pathophysiologic mechanisms involved in a murine model of the cardiometabolic HFpEF phenotype induced by high fat diet (HFD) and endothelial nitric oxide synthase (NOS) inhibitor (L-NAME) (21-23).

## Results

Average daily food and water intakes, estimated over 9-week HFD+L-NAME regimen **(Table S1)**, were markedly reduced in HFpEF mice (p<0.001 and p<0.0001, respectively) compared to control (normal water and normal diet), consistent with previous reports (24, 25); however, no differences were observed between HFpEF-placebo and HFpEF-MR-356 groups.

### Systemic and Cardiac Changes in HFpEF

The HFpEF phenotype developed following 5 weeks of HFD+L-NAME, confirmed with echocardiography **(Fig. S1A-D)**. At that stage, mice exhibited impaired diastolic function as evidenced by prolonged isovolumetric relaxation time [IVRT] (p<0.001), increased E/A ratio (p<0.05) and E/E’ ratio (p<0.0001) compared to controls. Ejection fraction (EF) was preserved (p=ns). HFpEF mice also developed hypertension (p<0.0001) and glucose intolerance (p<0.05) **(Fig. S1E-G)**. Together these results document cardiometabolic HFpEF, as previously described (23).

### GHRH-Agonist MR-356 counteracts the cardiometabolic HFpEF phenotype

We hypothesized that GHRH-Agonist MR-356 would reverse this phenotype and after 5-weeks of HFD-L-NAME diet mice were randomized to receive 4 weeks of daily injections of either placebo or MR-356. **Table S2** depicts parameters of body morphology [body weight (BW), tibia length (TL)], organ weight and lung water with and without normalization to body morphology after a 9-week regimen. As expected, BW **(Fig. 1A)** was substantially increased in HFD+L-NAME-fed mice in comparison to the control group (p<0.01). As shown in **Fig. 1B-C**, treatment with MR-356 tended to reduce the HW/TL ratio (p=0.055) and lung water content (LWC, p=0.051) in comparison to placebo-treated mice, suggesting the attenuation of cardiac hypertrophy and pulmonary congestion. Systolic (SBP, p<0.0001) and diastolic (DBP, p<0.001) blood pressure was similarly increased in both HFpEF groups compared to control mice **(Fig. 1D-E, Table S3)**. Intraperitoneal glucose tolerance test (ipGTT) revealed increased glucose levels in HFpEF-placebo mice versus control mice (p<0.05), suggesting impaired glucose tolerance **(Fig. 1F-G)**. Treatment with MR-356 eliminated the increased glucose levels following ipGTT in HFpEF vs. controls, except for the 15 min. time point (Fig 1G). Importantly, the cardiac phenotype after 9 weeks of the HFD+L-NAME regimen did not show differences in EF (p=ns) among groups **(Fig. 2A)** but the E/E’ ratio (p<0.01, **Fig. 2B**), global longitudinal strain (GLS, **Fig. 2C**) and exercise tolerance **(Fig. 2D)**, were all improved in mice treated with MR-356. Additional echocardiographic parameters are provided in **Table S4**. Representative echocardiographic images of M-mode tracings of parasternal short axis views and images of mitral inflow velocities by pulsed-wave Doppler and mitral annulus velocity by tissue Doppler imaging (TDI) are shown in **Fig S2**.

**Figure 1.**
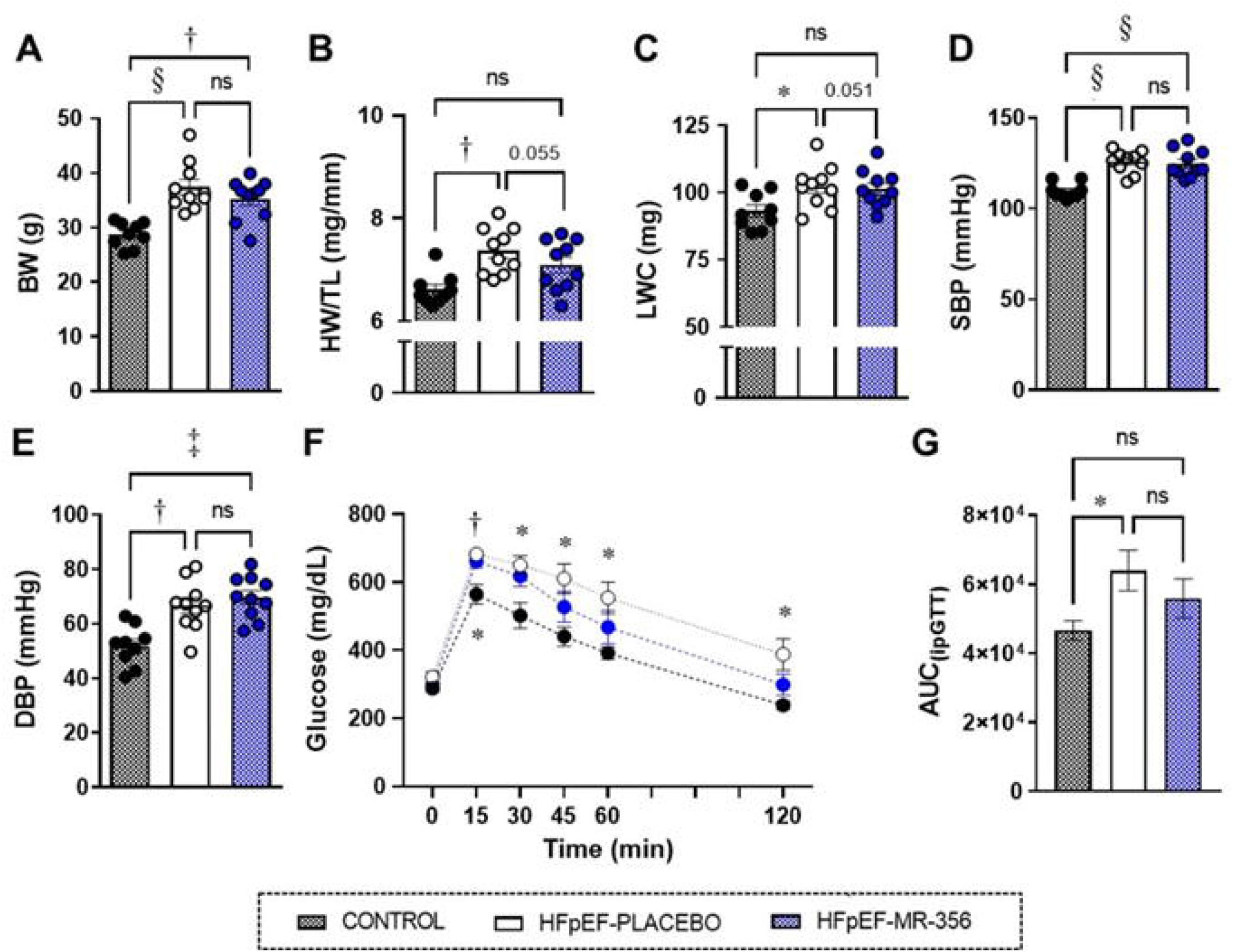
HFpEF phenotype after 9 weeks of HFD-L-NAME diet. **(A)** Body weight (BW) was substantially increased in HFD + L-NAME fed mice in comparison to control group. Treatment with MR-356 reduced the **(B)** heart weight (HW) to tibia length (TL) ratio (HW/TL) (p=0.055) and **(C)** lung water content (LWC) (p=0.051) in comparison to placebo-treated mice suggesting attenuation of cardiac hypertrophy and pulmonary congestion, respectively. **(D)** Systolic (SBP) and **(E)** diastolic (DBP) blood pressure values were substantially increased to a comparable extent in both HFpEF groups compared to control mice. All data are expressed as mean ± SEM. (One-way ANOVA followed by Tukey’s multiple comparisons test, * p<0.05, † p<0.01, ‡ p<0.001, § p<0.0001 vs. Control (Control: n=9, HFpEF-placebo: n=10 and HFpEF-MR-356: n=10). **(F)** Intraperitoneal glucose tolerance test (ipGTT) showed increased glucose levels in HFpEF-placebo mice compared to control mice in all time points after glucose injection while MR-356 treated mice showed an increase in glucose levels only at 15-minutes time point (Two-way ANOVA followed by Tukey’s multiple comparisons test, * p<0.05, † p<0.01 vs. Control, n=10/group). **(G)** The area under the curve (AUC) of ipGTT was increased in HFpEF-placebo group, suggesting impaired glucose tolerance. In contrast, MR-356 did not significantly attenuate glucose levels (One-way ANOVA followed by Tukey’s multiple comparisons test, * p<0.05 vs. control, n=10/group).

**Figure 2.**
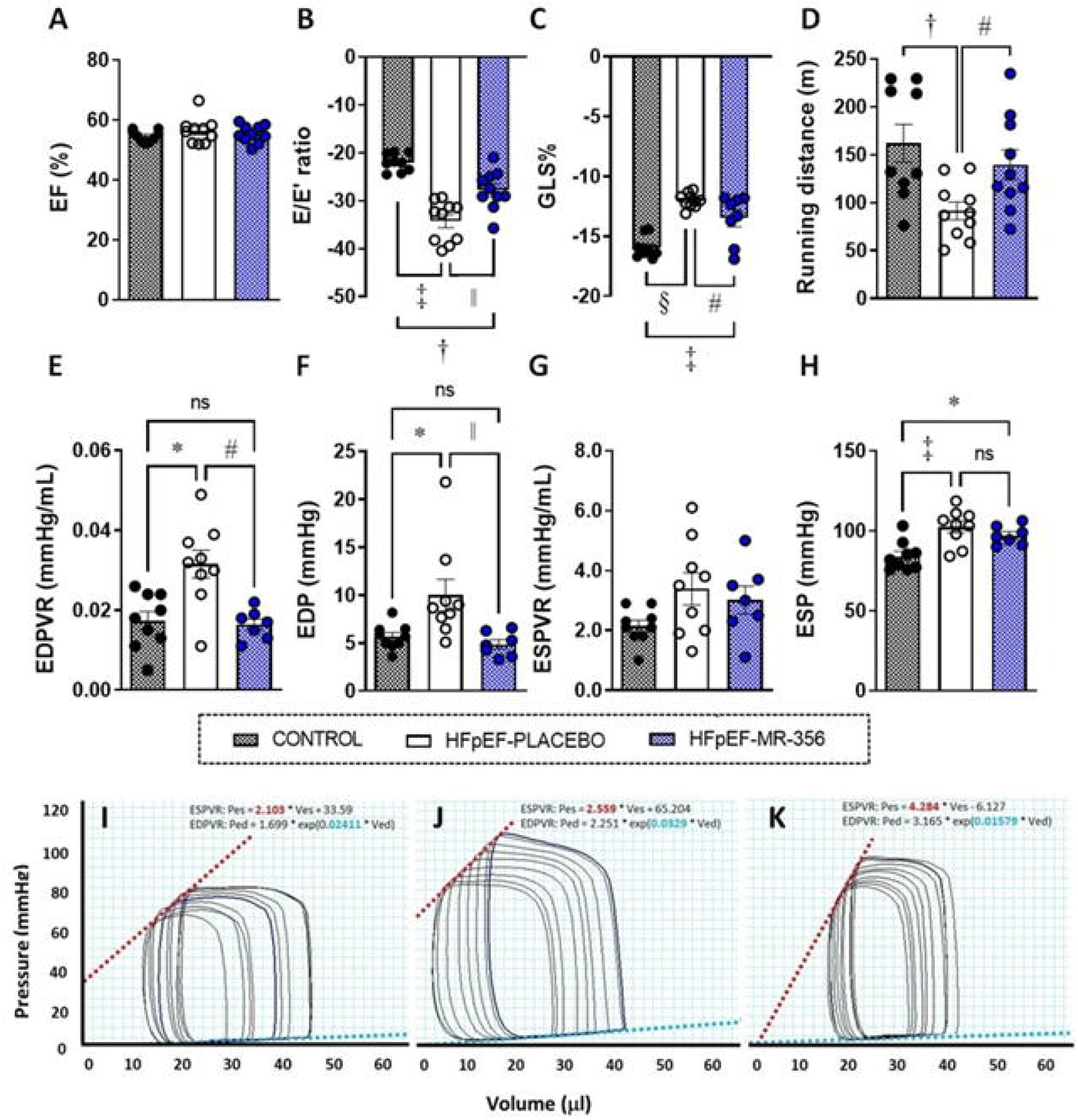
Cardiac performance and hemodynamic changes after 9 weeks of HFD+L-NAME diet. **(A)** Ejection fraction (EF) was not different among the experimental groups (p=ns); **(B)** the ratio between early mitral inflow velocity (E) and mitral annular early diastolic velocity (E’) [E/E’] was markedly increased (absolute values) in HFpEF mice in comparison to control mice while treatment with MR-356 improve diastolic dysfunction **(C)** global longitudinal strain (GLS) was reduced (absolute values) revealing impaired cardiac performance. MR-356 treatment significantly improved these parameters (One-Way ANOVA followed by a post hoc Holm-Sidak’s test, # p<0.05 and || p<0.01 vs. HFpEF-placebo, † p<0.01, ‡ p<0.001 and § p<0.0001 vs. Control, n=9-10). **(D)** Exercise tolerance test showed a reduced running capacity in the HFpEF-placebo mice compared to control and HFpEF-MR-356 mice. (One-Way ANOVA followed by a post hoc Holm-Sidak’s test, # p<0.05, † p<0.01, n=9-10). **(E)** Slope of end-diastolic pressure-volume relationship (EDPVR), **(F)** LV end-diastolic pressure (EDP), **(G)** slope of end-systolic pressure-volume relationship (ESPVR), and **(H)** LV end-systolic pressure (ESP). E-F: Kruskal-Wallis test followed by Dunn’s multiple comparisons test * p<0.05 and † p<0.01 vs. Control,, # p<0.05 and || p<0.01 vs. HFpEF-placebo, n=9-10. C-D: One-Way ANOVA with Tukey’s multiple comparisons test, * p<0.05, ‡ p<0.001, n=9-10. Bottom panels depict representative pressure-volume (PV) loops from control **(I)**, HFpEF-placebo **(J)** and HFpEF-MR-356 **(K)** during inferior vena cava occlusion. Red dashed lines represent ESPVR, and light teal dashed lines represent EDPVR. HFpEF-placebo mice show increased EDPVR and EDP indicating increased chamber stiffness. Notably, treatment with MR-356 restored these parameters to normal levels.

### Cardiovascular Hemodynamics

Heart rate (HR) values did not differ between control and HFD+L-NAME-fed mice. Consistent with echocardiography, abnormal diastolic function was evident from LV pressure-volume (PV) recordings in HFpEF-placebo mice **(Fig. 2E-F)** evidenced by steeper end-diastolic pressure-volume relationship (EDPVR, p<0.05) and elevated end-diastolic pressure (EDP, p<0.05) in comparison to control mice. Notably, treatment with MR-356 completely restored both diastolic parameters (p<0.05 and p<0.01, respectively vs. HFpEF-placebo group). LV end-systolic blood pressure (ESP), increased in HFD+L-NAME-fed mice, was not different between placebo- and MR-356-treated mice **(Fig. 2G)**. Similarly, there was no difference in the slope of end-diastolic pressure-volume relationship (ESPVR, p=ns) among the experimental groups. Additional hemodynamic parameters are given in **Table S5**.

### Cardiac Hypertrophy, Fibrosis and Capillary Density

HFpEF is associated with cardiac hypertrophy and increased fibrosis (26). We observed that both myocardial cross-sectional area (CSA, **Fig. 3A**) and cardiac interstitial fibrosis **(Fig. 3B)** were markedly increased in HFpEF-placebo mice (p<0.01) in comparison to controls whereas treatment with MR-356 completely restored these parameters (p<0.01). Similar to previous reports (22), capillary density **(Fig. 3C)** was substantially reduced in HFpEF-placebo mice in comparison to controls (p<0.01) whereas treatment with MR-356 attenuated capillary rarefaction (p<0.05 vs. HFpEF-placebo).

**Figure 3.**
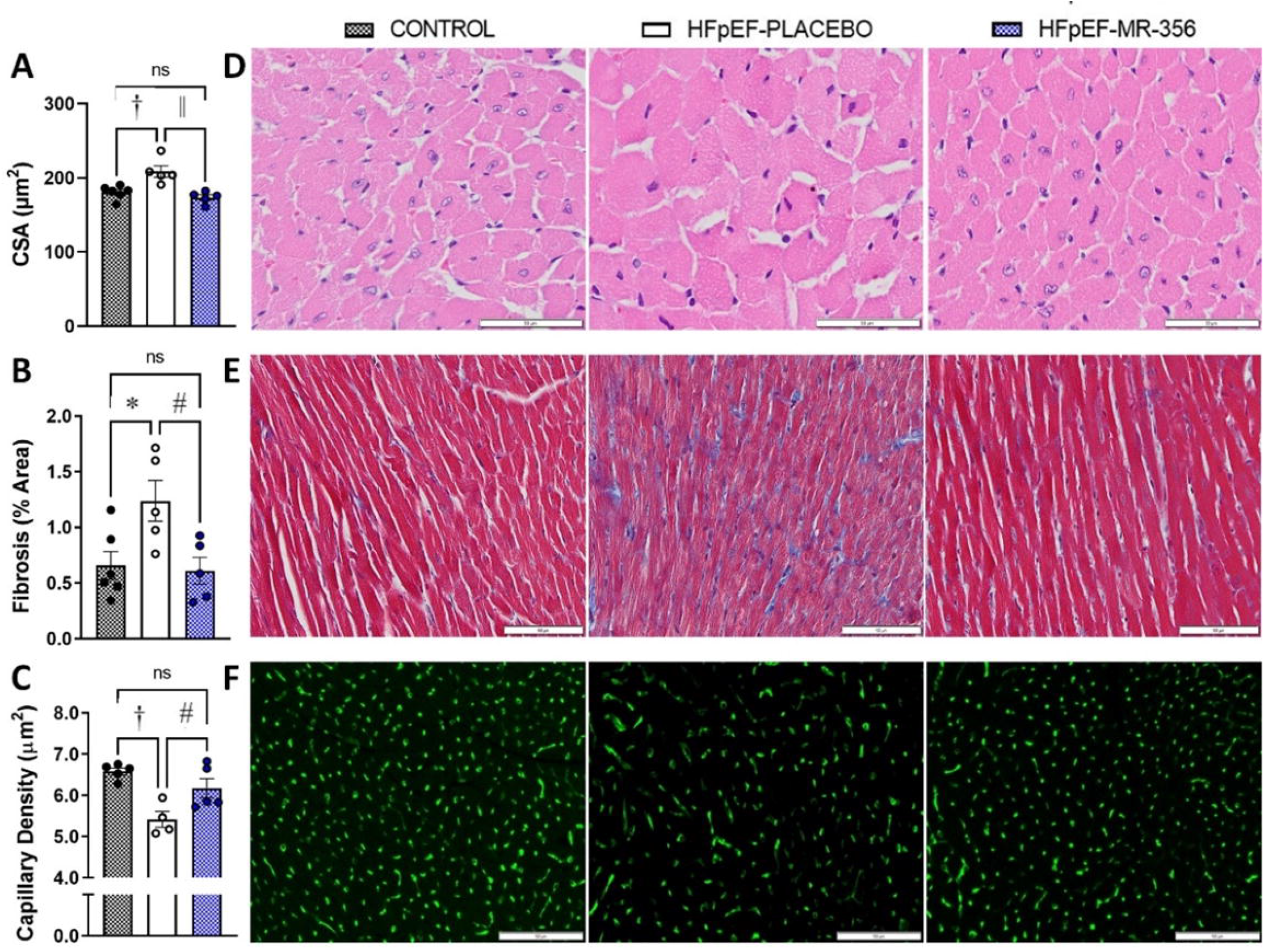
Cardiac hypertrophy, fibrosis, and capillary density. Bar graphs showing **(A)** increased myocardial cross-sectional area (CSA), **(B)** increased fibrosis and **(C)** capillary rarefaction in HFpEF-placebo mice in comparison to control and HFpEF-MR-356. One-Way ANOVA followed by Tukey’s test * p<0.05, † p<0.01 vs. Control, # p<0.05, || p<0.01 vs. HFpEF-placebo, n=4-6). Representative **(D)** hematoxylin and eosin (scale bar: 50 μm), **(E)** Masson’s trichrome (scale bar:100 μm) and **(F)** isolectin B4 (scale bar: 100 μm) stained sections from control mice **(left panels)**, HpEF-placebo **(middle panels)** and HFpEF-MR-356 **(right panels)**, respectively.

### Pro-BNP, iNOS and VEGF-A Expression in HFpEF

Elevated levels of pro-BNP are a characteristic of HFpEF in humans (27). Hearts of HFpEF-placebo mice exhibited significantly increased pro-BNP expression **(Fig. 4A, I)** compared to control mice (p<0.05), while the level of pro-BNP was restored to normal in mice treated with MR-356 (p<0.01). Next, we investigated the expression of inducible nitric oxide synthase (iNOS) and vascular endothelial growth factor A (VEGF-A). Several studies demonstrate that iNOS is minimally expressed in normal hearts; however, upon stress, iNOS is activated and induces oxidative stress and mitochondrial dysfunction in CMs (28, 29), both of which contribute to the development of HFpEF observed in experimental and clinical settings (23). Elevated levels of VEGF-A are also associated with cardiac hypertrophy and impaired cardiac performance (30). HFpEF-placebo mice exhibited an upregulation of iNOS expression **(Fig. 4B, J)** compared to the control group (p<0.05) while no such increase was noted in HFpEF-MR-356 mice (p=ns). In fact, iNOS expression in the HFpEF-MR-356 group was reduced to control levels (p<0.05 vs. HFpEF-placebo). Similarly, expression of VEGF-A **(Fig. 4C, K)**, markedly increased in the HFpEF-placebo group compared to the control group (p<0.001), was restored to normal levels by treatment with MR-356 (p<0.05). The increase in myocardial CSA, indicative of cardiac hypertrophy is consistent with the changes in the expression of VEGFA observed here.

**Figure. 4.**
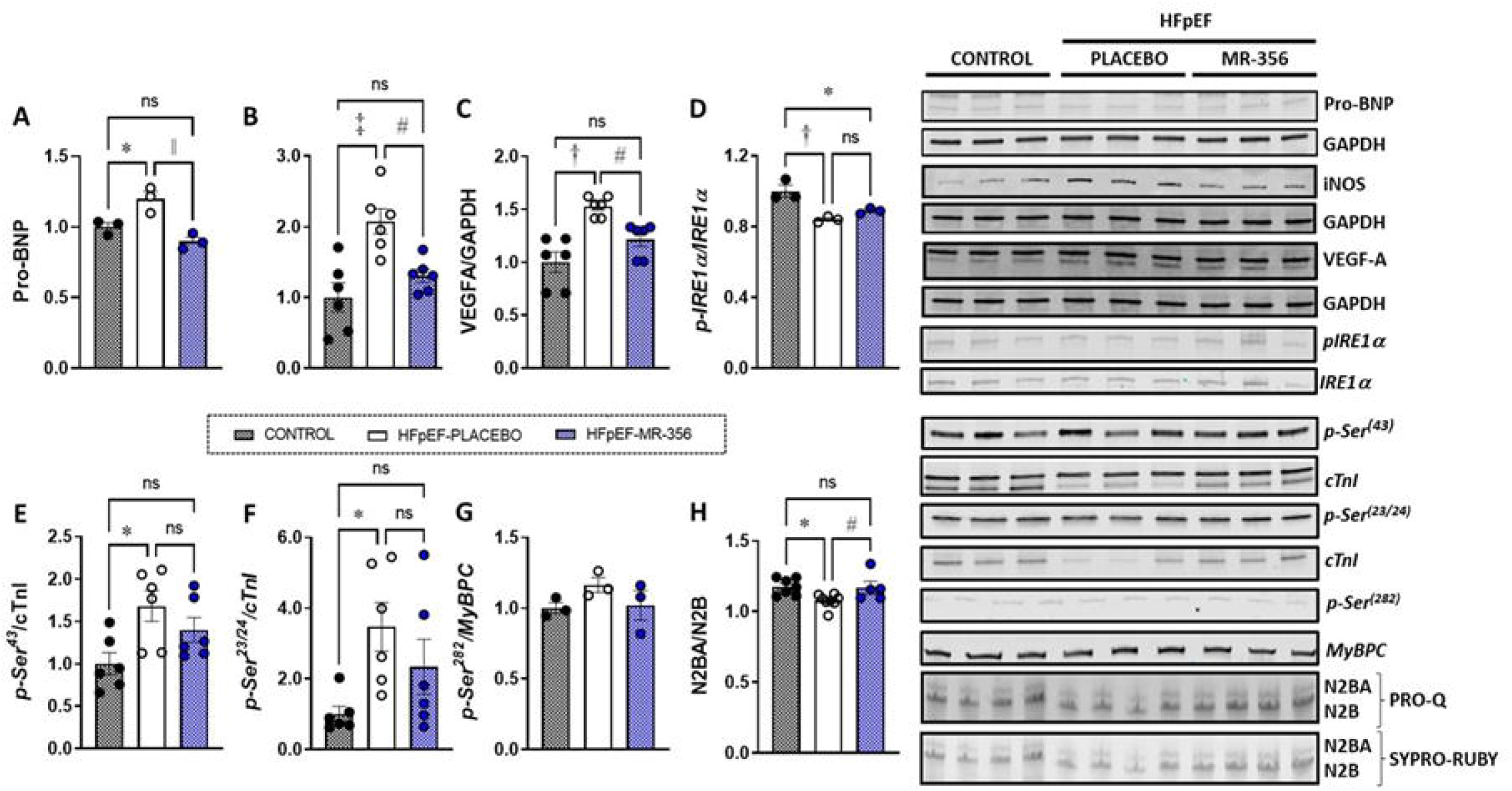
Changes in cardiac expression of pro-BNP, iNOS and VEGFA in HFpEF mice. Bar graphs showing increased expression of cardiac pro-BNP **(A)**, iNOS **(B)** and VEGFA **(C)** expression in HFpEF-placebo mice in comparison to control and HFpEF-GHRH-A mice (One-way ANOVA followed by Tukey’s test * p<0.05, † p<0.01 vs. Control, # p<0.05, || p<0.01 vs. HFpEF-placebo, n=6). **(D)** Western blot quantification showing reduced ratio of phospho-IRE1α/IRE1α in HFpEF-placebo mice in comparison to control (One-Way ANOVA followed by Tukey’s multiple comparisons’ test, * p<0.05, † p<0.01 vs. Control, n=3). Western blot analysis of expression of **(E)** phosphorylation of cTnI at Ser^43^ (p-Ser^43^) (One-way ANOVA followed by Tukey’s multiple comparisons test, * p<0.05 vs. control, n=6) and **(F)** phosphorylation of cTnI at Ser^23/24^ (p-Ser^23/24^) (Kruskal-Wallis test followed by Dunn’s multiple comparisons test, * p<0.05 vs. Control, n=6) normalized to the total cTnI abundance in LV homogenates. **(G)** Phosphorylation of myosin binding protein C (MyBPC) at Ser^282^ (p-Ser^282^) expression normalized to the total MyBPC abundance in LV homogenates. **(H)** Bar graphs show titin isoform expression from control (n=7), HFpEF-placebo (n=8) and HFpEF-MR-356 (n=5) mice (One-way ANOVA followed by Newman-Keuls’ multiple comparisons test, * p<0.05 vs. Control, # p<0.05 vs. HFpEF-placebo). Representative immunoblots **(I-K)** of pro-BNP, iNOS, VEGFA and respective loading control (GAPDH) for each experimental group (n=3); and **(L)** immunoblots of phosphorylated IRE1α at Ser^724^ and total IRE1α from heart samples (n=3) of each experimental group. **(M & N)** Phosphorylation status of cTnI (Ser^43^ and Ser^23/24^, respectively) and total cTnI as well as phosphorylation of MyBPC (Ser^282^) and total MyBPC **(O)**. Representative 1% agarose gel **(P)** showing total titin phosphorylation using Pro-Q Diamond staining and SYPRO-RUBY staining for total titin. Titin N2BA corresponds to the more compliant isoform (upper band) while N2B corresponds to the stiffer isoform (lower band).

### Effect of GHRH-Agonist MR-356 on Regulation of Cardiomyocyte (CM) Stress Responsiveness

We next examined the expression of inositol-requiring enzyme 1α (IRE1α) in hearts from all three groups of mice to evaluate the ability of MR-356 to modulate the unfolded protein response (UPR) (**Fig. 4D, L)**. The IRE1α-X-box-binding protein 1(XBP1) axis, part of the UPR is a regulator of CM stress responsiveness is dysregulated in experimental and clinical HFpEF (23). In line with previous studies, both HFpEF groups showed reduced expression of phosphorylated IRE1α as compared to controls (p<0.05). MR-356 had no apparent effect in restoring this pathway.

### Phosphorylation of Sarcomeric Proteins

Phosphorylation of cardiac contractile proteins, particularly, cardiac troponin I (cTnI) and myosin binding protein C (MyBPC) play important roles in the modulation of sarcomeric function, particularly, on cardiac contractility and pump function (31, 32). The phosphorylation status of cTnI at Ser43 **(Fig. 4E, M)** and Ser 23/24 **(Fig. 4F, N)** was increased in HFpEF-placebo mice (p<0.05) whereas treatment with MR-356 appeared to reduce these levels, as HFpEF-MR-356 mice exhibited no significant difference compared to control levels. Phosphorylation of MyBPC can alter the sensitivity of the myofilaments to calcium and, thus, contribute to modulation of the CM stiffness. Contrary to cTnI, there was no difference in phosphorylation of MyBPC between the control and experimental groups **(Fig. 4G, O)**.

### Titin Modifications in HFpEF

The giant sarcomeric protein titin, along with collagen, is the main determinant of CM stiffness (33). Both short-term post-translational modifications of titin and changes in the ratio of the N2BA/N2B isoforms in the long-term can mediate increased CM stiffness. Consistent with previous reports (34, 35) and our findings in the porcine model CKD-induced HFpEF (12), our data showed hypophosphorylation of titin as the ratio of phosphorylated N2BA/N2B was markedly reduced in HFpEF-placebo mice (p<0.05) which was attenuated by treatment with MR-356 **(Fig. 4H, P)**.

## Discussion

Our findings demonstrate that administration of the GHRH agonist, MR-356, improved most of the key phenotypes associated with a murine model of cardiometabolic HFpEF (22, 23). In this study, we recapitulated the cardiac and systemic changes associated with the cardiometabolic HFpEF phenotype by feeding mice a HFD (metabolic stress) and administering a NOS inhibitor (L-NAME), which together result in an increase in body weight, systolic and diastolic pressure, and impaired glucose and exercise tolerance (21-23). In addition, these mice developed cardiac hypertrophy, fibrosis, and impaired diastolic function while EF was preserved. Diastolic dysfunction was supported by invasive hemodynamic measurements showing increased EDPVR and EDP which characterizes increased ventricular chamber stiffness (36). Notably, our findings revealed restoration towards normal of exercise capacity in mice treated with MR-356, consistent with recent findings in aged mice treated with agonist of GHRH MR-409 (37).

Our findings are in agreement with earlier studies showing salutary effects of GHRH-agonists in cardiorenal models of HFpEF (11, 12) and in reversing cardiac hypertrophy due to aortic banding (10). The present findings are an important advance, addressing a crucial underpinning of HFpEF due to cardiometabolic derangements.

### GHRH-Agonist MR-356 Reduces Myocardial Wall Stress in HFpEF

Several biochemical measurements, including pro-BNP and immune activation, support the underlying favorable effects of GHRH-A on the myocardium. First, GHRH-A restored elevated pro-BNP levels toward normal. Pro-BNP participates in CV homeostasis through diuretic, natriuretic, vasorelaxant, anti-proliferative, anti-inflammatory, and anti-hypertrophic effects (38, 39), and is an effective marker of myocyte stress, correlating with cardiac hypertrophy and fibrosis (40). This result is supported by our hemodynamic findings demonstrating a beneficial effect of MR-356 treatment on the slope of EDPVR. Reduction in the pro-BNP levels was similarly noted in our porcine CKD-HFpEF model (12).

### Effect of GHRH-Agonist MR-356 on Inflammation

Second, GHRH-A restored several markers of myocardial inflammation to normal. Endothelial dysfunction has been proposed to link chronic systemic inflammation with myocardial dysfunction and remodeling in HF (41-43). Schiattarella *et al*. (23) suggested a role of iNOS activation and metabolic inflammation in HFpEF pathogenesis. VEGF-A is also produced by cardiomyocytes during inflammation, mechanical stress, and cytokine stimulation (44), and elevated VEGF-A expression is often associated with poor prognosis and disease severity (30, 44). In line with these findings, we showed that MR-356 downregulated the expression of iNOS and VEGFA in the heart. VEGF-A-induced cardiac hypertrophy is associated with increased Akt signaling (30). Our results showed an increase in VEGF-A and pro-BNP expression in the HFpEF-placebo hearts but no changes in phosphorylation of Akt; however, our study was based on phosphorylation of Akt at threonine 308 (Thr308) and not at Ser473. Similar to a previous report (23), we observed a suppression of IRE-1α phosphorylation in HFpEF mice although IRE-1α levels were not affected by MR-356.

### GHRH-Agonist MR-356 Reversed Cardiac Hypertrophy, Fibrosis and Capillary Rarefaction

HFpEF-MR-356 mice showed no evidence of cardiac hypertrophy, fibrosis, and capillary rarefaction. To assess signaling pathways relevant to hypertrophy and fibrosis and to test the possible involvement of several PKG-associated downstream targets in HFD+L-NAME-induced HFpEF, we analyzed the activation of ERK 1/2 and AKT in cardiac tissues. Surprisingly, quantitative immunoblotting showed that phosphorylated ERK 1/2 (Thr202/203) was not altered in the HFpEF hearts while phosphorylated Akt was modestly but not significantly increased in HFpEF-MR-356 mice **(Fig. S3)**.

### GHRH-Agonist MR-356 Improves Diastolic Function

Diastolic dysfunction has been associated with reduced phosphorylation of the myofilament regulatory proteins. In fact, phosphorylation of cTnI is a significant factor in cardiac contraction and relaxation (31, 32); however, the effects of cTnI phosphorylation depend upon the combination of kinases activated, the specific sites phosphorylated, and the activities of protein phosphatases (31). Site-specific phosphorylation of cTnI at Ser 23/24 by protein kinase A (PKA) or protein kinase G (PKG) is associated with reduced myofilament Ca^2+^ sensitivity and increased crossbridge cycling rate and enhanced unloaded shortening velocity, which also contribute to β-agonist-induced lusitropy. Conversely, phosphorylation at Ser43 by protein kinase C (PKC) is associated with reduced maximum Ca^2+^-activated force and decreased crossbridge cycling rates (31, 45), which are likely to delay relaxation with an increased economy of contraction (31). Our results showed increased phosphorylation at Ser23/24 and Ser43 in the placebo mice in comparison to control mice with no changes in phosphorylation of MyBPC at Ser282. These results differ from our recent findings in the murine model induced by chronic administration of low dose of angiotensin-II (Ang-II) (11). However, the HFD-L-NAME-fed mice were subjected to higher stress producing conditions (including metabolic challenge and physical activity) while the Ang-II-induced HFpEF mice received regular food and were mostly sedentary. This difference might have led to hyperactivation of the sympathetic system in the HFD-L-NAME groups, which could have produced elevated phosphorylation background (46) mediated by PKA. Therefore, the functional significance of the multiple phosphorylation sites of cTnI would strongly depend on the integration of the entire myofilament phosphorylation background in such conditions (47).

### Treatment with GHRH-Agonist MR-356 Restores Titin Phosphorylation Status

Hypophosphorylation of titin in the myocardium is considered a molecular hallmark of HFpEF (48) and is associated with reduced PKG activity resulting from oxidative stress which is consistent with our recent work showing an increased cGMP in CMs upon GHRH-MR-356 treatment in an AngII-induced HFpEF phenotype (12). Titin and collagen are the major determinants of myocardial stiffness (49) and titin’s contribution to myocardial stiffness can be modulated by isoform shift and phosphorylation status (50). We recently showed that a related GHRH-agonist, MR-409, improves the phosphorylation of titin isoforms in a porcine model of CKD-induced HFpEF (12). Similarly, our data support the contribution of MR-356 in restoring the phosphorylation of titin isoforms and, thus, reducing myocardial stiffness in this model. Notably in our current study, we assessed total titin phosphorylation using ProQ Diamond staining which can detect PKA phosphorylation of the N2B but does not specifically detect PKC phosphorylation which requires phospho-specific antibodies (51).

Currently, advances in therapeutic approaches to treat HFpEF are limited due mostly to poor understanding of the underlying mechanisms involved in the predominant HFpEF-phenotype. Given that HFpEF is a multifactorial disease and has heterogenous and complex clinical presentations (52), it is reasonable to assume that there may be multiple mechanisms involved; thus, the need for development of animal models that can mimic features of HFpEF-phenotypes is crucial for advancing HFpEF treatments. Notably, we reported beneficial effects of therapy with GHRH-agonist not only in HFrEF (6, 7, 13, 53) but also HFpEF in small (11) and large animal (12) models. Importantly, the beneficial effects of agonists of GHRH occurred by preventing the onset of HF (11) as well as in reversing established HFpEF (11, 12). The results of the present study indicate that treatment with the GHRH-agonist, MR-356, can block or attenuate the majority of the key features of HFpEF such as cardiac hypertrophy, diastolic dysfunction, fibrosis, capillary rarefaction, and myocardial stiffness, suggesting GHRH agonists may be a novel approach to targeting multiple organ system derangements observed in the HFpEF. Strikingly, our results are supported by evidence of the GHRH’s broad range of regulatory influences in the cardiovascular (CV) system and in aging (37), where these compounds produce therapeutic effects in several cardiometabolic conditions including obesity (15) and diabetes (17, 18).

Accordingly, our findings reveal that GHRH-agonist therapy improves a cardiometabolic HFpEF phenotype suggesting that activation of GHRH receptor signaling in the heart as a potential therapeutic strategy for treatment of HFpEF. The insights presented here provide support for ongoing translational development of this class of molecules for HFpEF.

## Materials and Methods

An extended version of Materials and Methods can be found in *SI Appendix, Supporting Information* (*SI) Material and Methods*.

The protocol of the resent study was reviewed and approved by the University of Miami Institutional Animal Care and Use Committee and complies with all Federal and State guidelines concerning the use of animals in research and teaching as defined by *The Guide for the Care and use of Laboratory Animals* (54) and the ARRIVE Guidelines (55). Animal Welfare Assurance # A-3224-01, approved through November 30, 2023. Protocol # 21-044 (Approval: 03/17/2021; Expiration: 03/16/2024). The University of Miami has received full accreditation with the Association for Assessment and Accreditation of Laboratory Animal Care (AAALAC International), site 001069.

### Animal model

C57BL6N (Charles River Laboratories) male mice (8-weeks-old) were fed a high-fat diet (HFD, D12492, Research Diets, Inc) plus the nitric oxide synthase (NOS) inhibitor, N^**ω**^ – nitro-L-arginine methyl ester (L-NAME, 0.5 g/L, Sigma Aldrich #N5751) for 9 weeks to induce HFpEF (HFD+L-NAME) (23). Five weeks after beginning the HFD+L-NAME regimen, mice were randomly assigned to receive daily subcutaneous injections (100 μl) of either vehicle (DMSO + propylene glycol, HFpEF-placebo) or GHRH-agonist MR-356 (HFpEF-MR-356, 200 μg/Kg/day) for 4 weeks. Control animals received regular chow and water. The timeline and brief description of the protocol are shown in **Fig S4**.

### Drugs

L-NAME (Sigma-Aldrich) was provided in the drinking water (0.5 g/L) ad-libitum. MR-356 was synthesized and purified in the laboratory of one of us (A.V.S.) by Ren-Zhi Cai and Wei Sha. The peptide was dissolved in an aqueous solution of 0.1% DMSO (Sigma) and 10% propylene glycol (Sigma) as previously described (7).

### Blood Pressure

Blood pressure was measured noninvasively in conscious mice using the tail-cuff BP-2000 Blood Pressure Analysis System™ (Visitech System, Inc). Mice were acclimated in individual restrain holders on a temperature-controlled platform (37°C), and blood pressure recordings were performed at the same time of the day for 3-5 consecutive days. Readings from the last session was averaged from at least 12-15 measurements.

### Intraperitoneal Glucose Tolerance Test (ipGTT)

Intraperitoneal glucose tolerance tests were performed by injection of glucose (2 g/kg in saline) after 6-hour fasting in a quiet and stress-free environment at 5- and 9-week timepoints. Tail blood glucose levels (mg/dL) were measured with a glucometer (AlphaTrak II, Zoeltis) before (0 min) and at 15, 30, 45, 60, and 120 minutes after glucose administration, and the area under the curve (AUC) was calculated.

### Exercise Tolerance Test (EET)

Exercise protocol followed the American Physiological Society’s Resource Book for the Design of Animal Exercise Protocols guidelines (56). Mice were acclimated for 3 days to treadmill exercise; an exhaustion test was performed as previously reported with minor modifications (23).

### Echocardiography

Serial cardiac images were acquired using the Vevo^®^2100 imaging system (FUJIFILM VisualSonics Inc.) equipped with a high-resolution transducer (MS400) to characterize the HFpEF phenotype as previously described (11). Briefly, mice were anesthetized with 1-3% isoflurane and transferred to a platform where body temperature and heart rate (HR) were monitored throughout the examination. Two-dimensional (B-mode) and M-mode images from the parasternal long-axis and short-axis (at the papillary muscle level), and four-chamber apical views were acquired during the exam for cardiac morphology and function. Echocardiographic parameters were analyzed by standard and speckle tracking echocardiography (STE) analysis off-line using Vevo Lab software (version). Relative wall thickness in diastole (RWTd) was calculated by the following formula: RWTd=2xPWd/LVEDD while LV remodeling index (LVRI) was calculated as LV mass/LVEDD. All B-mode and M-mode measurements were obtained using AutoLV analysis software to minimize beat to beat variability and bias. Pulsed-waved Doppler and tissue Doppler measurements of the mitral valve were obtained from the apical four chamber view. For speckle tracking echocardiography (STE) B-mode images were obtained from the parasternal longitudinal axis view, three consecutive cardiac cycles were selected for analysis and semi-automated tracing of the endocardial and epicardial border were obtained to determine global longitudinal strain (GLS).

### Pressure-Volume (PV) Loops

Hemodynamic studies were performed using a micro-tipped pressure-volume catheter (SPR-839; Millar Instruments) as previously described with minor modifications (11, 57). Briefly, mice were induced and maintained with isoflurane, and body temperature was controlled (∼37°C) during the whole procedure. The left internal jugular vein was exposed and cannulated with a 30-gauge needle for the administration of fluid support. The right carotid artery was exposed to permit the catheter to advance into the LV. Pressure-volume (PV) loops were recorded during steady-state and temporary inferior vena cava (ivc) occlusion. The ventilator was stopped momentarily (∼10 s) to avoid breathing interference during measurements. The volumes were calibrated using volumes derived from echocardiographic measurements. All analyses were performed using LabChart Pro version 8.1.5 software (ADInstruments).

### Post-mortem analysis

After hemodynamic studies, blood was withdrawn, the animals (under deep anesthesia) were humanely euthanized, and their tissues were harvested for further analysis. The whole heart and lungs were weighed, and tibia length was measured for normalization. Next, pulmonary congestion was evaluated by assessing the lung water content (LWC) which was calculated by subtracting lung dry weight from the lung wet weight measured immediately after excision. Lungs were dried (∼72 hours) in an oven at 37°C (58).

### Histology

Heart sections were scanned using a whole slide digital microscopy imaging system (Olympus VS120 Scanner, OlyVIA 2.9 software) to measure cardiomyocyte cross sectional area (CSA), cardiac fibrosis and capillary density as previously described with minor modifications (11). All measurements were performed using Image J by an investigator masked to treatment assignment.

### Western Blotting

Flash-frozen cardiac samples were homogenized using a Pyrex glass-glass homogenizer on liquid nitrogen and urea buffer containing: 8 M urea, 2 M thiourea, 3% SDS, 0.03% bromophenol blue and 0.05 M Tris, pH 6.8. Glycerol (50%) with protease inhibitors ([in mmol/liter] 0.04 E64, 0.16 leupeptin, and 0.2 PMSF) at 60°C for 10 min was added. Samples were centrifuged at 12500 RPM for 5 minutes, aliquoted, flash frozen in liquid nitrogen, and stored at -80°C as previously described (11, 12). Immunoblot analysis was performed using antibodies listed in **Table S6**. Images were acquired by an Odyssey infrared imaging system (LI-COR Biosciences, Lincoln, NE) and then analyzed by the Image J software (NIH). Quantification of experiments are represented as fold change compared to control.

### Titin Analysis

For detection of titin, LV samples were solubilized and electrophoresed using 1% agarose gels as described previously (12). Total titin phosphorylation was analyzed using the fluorescence-based phosphoprotein stain ProQ diamond (Invitrogen) in comparison to the total protein stain Sypro® Ruby (Invitrogen).

### Statistical Analysis

Results are expressed as mean ± standard error of mean (SEM). All data were tested for Gaussian distribution using the Shapiro-Wilk test. Data between HFD+L-NAME ± GHRH-MR-356 treatment were compared using unpaired Student’s *t*-test or ANOVA followed by Tukey’s post-hoc when multiple groups were involved or otherwise stated. For data that were not normally distributed, we used non-parametric tests: two-tailed Mann-Whitney U-test or Wilcoxon test while Kruskall-Wallis followed by Dunn’s multiple comparisons test was used to compare more than two groups. Differences in a series of time points were compared using two-way ANOVA for repeated measurements (RM). For a given parameter, p<0.05 was considered significant. All data analyses and plots were generated using GraphPad Prism 9 version 9.4.0 (GraphPad Software Inc. CA, USA). Symbols represent p-values of different orders of magnitude, *p<0.05, †p< 0.01, ‡p<0.001, §p<0.0001 and ¶p<0.00001 vs. Control; and #p<0.05, ||p<0.01 vs. HFpEF-Placebo.

## Acknowledgments

We thank the Analytical Imaging Core Facility at the University of Miami Miller School of Medicine for the VS120 Slide Scanner (NIH grant 1S10OD023579-01).

