## Supplemental Information for "Efficacy of a Growth Hormone-Releasing Hormone Agonist in a Murine Model of Cardiometabolic Heart Failure with Preserved Ejection Fraction"

***Exercise Tolerance Test (EET).*** Exercise protocol followed the American Physiological Society's Resource Book for the Design of Animal Exercise Protocols guidelines (5). Mice were acclimated for 3 days to treadmill exercise; an exhaustion test was performed as previously reported with minor modifications (3). Briefly, on the first day of training mice ran on the treadmill (Columbus Instruments) starting at a warm-up speed of 1.5 to 3.0 m/min, then the treadmill speed was slowly increased to 8 m/min for ~5 min after which speed was increased to 9 m/min at 5 min, 10 m/min at 7 min, and the treadmill was stopped at 10 min. On the second and third day of training, mice ran starting at a warm-up speed of 10 m/min, the treadmill speed was then increased to 11 m/min at 5 min, 12 m/min at 10 min, and the treadmill was stopped at 15 min. Mice were allowed to rest for 24 hours before the fatigue test. On the fifth day animals ran on a treadmill starting at a warm-up speed of 5 m/min for 4 min after which the speed was increased to 14 m/min for 2 min. Every subsequent 2 minutes, the speed was increased by 2 m/min until the animal was unable to return to running within 10 s of direct contact with an electric-stimulus grid. Running time and running distance were recorded.

***Echocardiography.*** Serial cardiac images were acquired using the Vevo®2100 imaging system (FUJIFILM VisualSonics Inc.) equipped with a high-resolution transducer (MS400) to characterize the HFpEF phenotype as previously described (6). Briefly, mice were anesthetized with 1-3% isoflurane and transferred to a platform where body temperature and heart rate (HR) were monitored throughout the examination. Two-dimensional (B-mode) and M-mode images from the parasternal long-axis and short-axis (at the papillary muscle level), and four-chamber apical views were acquired during the exam for cardiac morphology and function. Echocardiographic parameters were analyzed by conventional and speckle tracking echocardiography (STE) analysis off-line using Vevo Lab software (version).

Left ventricle end-diastolic (LVEDD) and end-systolic (LVESD) dimensions, end-diastolic (EDV) and end-systolic (ESV) volumes, posterior (PWT) and anterior wall thickness (AWT) were determined. Relative wall thickness in diastole (RWTd) was calculated by the following formula: RWTd=2xPWd/LVEDD while LV remodeling index (LVRI) was calculated as LV mass/LVEDD. Systolic function was assessed by LV ejection fraction (EF) calculated from cine-images in the parasternal long-axis view . All B-mode and M-mode measurements were obtained using AutoLV analysis software to minimize beat to beat variability and bias. Pulsed-waved Doppler and tissue Doppler measurements of the mitral valve were obtained from the apical four chamber view. Early (E wave) and late diastolic peak velocity (A wave) of mitral inflow, isovolumetric relaxation time (IVRT) and isovolumetric contraction time (IVCT), and tissue Doppler (E' and A' waves) velocity traces of mitral valve annulus at the septal level were obtained to assess diastolic function. The E/A and E/E' ratios, and myocardial performance index (MPI) were then calculated. For speckle tracking echocardiography (STE) B-mode images were obtained from the parasternal longitudinal axis view, three consecutive cardiac cycles were selected for analysis and semi-automated tracing of the endocardial and epicardial border were obtained to determine global longitudinal strain (GLS).

***Pressure-Volume (PV) Loops.*** Hemodynamic studies were performed using a micro-tipped pressure-volume catheter (SPR-839; Millar Instruments) as previously described with minor modifications (6, 7). Briefly, mice were induced and maintained with isoflurane, and body temperature was controlled (~37°C) during the whole procedure. The left internal jugular vein was exposed and cannulated with a 30-gauge needle for the administration of fluid support. The right carotid artery was exposed to permit the catheter to advance into the LV. Pressure-volume (PV) loops were recorded during steady-state and temporary inferior vena cava (ivc) occlusion. Occlusions of the ivc were performed by pushing the belly with a cotton tip or by direct occlusion using a small forceps after opening the chest (~4-5th intercostal space). The ventilator was stopped momentarily (~10 s) to avoid breathing interference during measurements. The volumes were calibrated using volumes derived from echocardiographic measurements. All analyses were performed using LabChart Pro version 8.1.5 software (ADInstruments).

***Post-mortem analysis.*** After hemodynamic studies, blood was withdrawn, the animals (under deep anesthesia) were humanely euthanized, and their tissues were harvested for further analysis. The whole heart and lungs were weighed, and tibia length was measured for normalization. Next, pulmonary congestion was evaluated by assessing the lung water content (LWC) which was calculated by subtracting lung dry weight from the lung wet weight measured immediately after excision. Lungs were dried (~72 hours) in an oven at 37°C (8).

***Histology.*** Heart sections were scanned using a whole slide digital microscopy imaging system (Olympus VS120 Scanner, OlyVIA 2.9 software) to measure cardiomyocyte cross sectional area (CSA), cardiac fibrosis and capillary density as previously described with minor modifications (6). All measurements were performed using Image J by an investigator masked to treatment assignment.

***Cardiac hypertrophy:*** Slides stained with hematoxylin and eosin (H&E) were used to measure cardiomyocyte (CM) cross-sectional area (CSA). Briefly, transverse sections of the myocardium were measured in 5-6 randomly high-power fields. We measured the area of approximately 300 CMs per heart section.

***Cardiac fibrosis:*** Heart sections stained with Masson's trichrome were used to quantify area of fibrosis. The relative amount of fibrosis area to total tissue was measured in each image using a color threshold technique.

***Capillary Density:*** To assess capillary density in the myocardium, heart sections were stained with isolectin B4 (GS-IB4-Alexa Fluor™488 conjugate, Invitrogen). Capillary density was quantified by dividing the number of isolectin B4+ cells by the area of randomly selected sections. Note that a size criterion of 10 μm was used to exclude small vessels. Each data point represents an average of five sections (at 20 x magnification) measured per heart.

***Western Blotting.*** Flash-frozen cardiac samples were homogenized using a Pyrex glass-glass homogenizer on liquid nitrogen and urea buffer containing: 8 M urea, 2 M thiourea, 3% SDS, 0.03% bromophenol blue and 0.05 M Tris, pH 6.8. Glycerol (50%) with protease inhibitors ([in mmol/liter] 0.04 E64, 0.16 leupeptin, and 0.2 PMSF) at 60°C for 10 min was added. Samples were centrifuged at 12500 RPM for 5 minutes. Then supernatants were collected and sampled for protein concentration by Bradford assay (Thermo Scientific), aliquoted, flash frozen in liquid nitrogen, and stored at -80°C. Samples (18 μg of protein) and dual molecular weight ladders (Precision Plus™ Protein Dual Color Standards, BioRad) were loaded on a 4-20% gradient agarose gel, electrophoresed, and transferred to nitrocellulose membranes (Bio-Rad Laboratories) as convenient. Membranes were blocked with 1% Tween-TBS + Rockland blocking buffer MB-070 (1:1). Immunoblot analysis was performed using antibodies listed in **Table S6**. Images were acquired by an Odyssey infrared imaging system (LI-COR Biosciences, Lincoln, NE) and then analyzed by the Image J software (NIH). Quantification of experiments are represented as fold change compared to control.

***Titin Analysis.*** For detection of titin, LV samples were solubilized and electrophoresed using 1% agarose gels as described with minor modifications (9). Total titin phosphorylation was analyzed using the fluorescence-based phosphoprotein stain ProQ diamond (Invitrogen) in comparison to the total protein stain Sypro® Ruby (Invitrogen). The gels were stained for 60 minutes with ProQ Diamond (Molecular Probes), washed and subsequently stained with SYPRO® Ruby (Molecular Probes). Images were scanned using a commercial scanner (Amersham Imager 680, GE corporation), and the intensities of the bands were quantified using Image J software as previously described (9). Titin isoform expression was determined in LV samples of control, HFpEF placebo and HFpEF-MR356 mice(n=6/group).

***Statistical Analysis.*** Results are expressed as mean ± standard error of mean (SEM). All data were tested for Gaussian distribution using the Shapiro-Wilk test. Data between HFD+L-NAME ± GHRH-MR-356 treatment were compared using unpaired Student's *t*-test or ANOVA followed by Tukey's post-hoc when multiple groups were involved or otherwise stated. For data that were not normally distributed, we used non-parametric tests: two-tailed Mann-Whitney U-test or Wilcoxon test while Kruskall-Wallis followed by Dunn's multiple comparisons test was used to compare more than two groups. Differences in a series of time points were compared using two-way ANOVA for repeated measurements (RM). For a given parameter, $p<0.05$ was considered significant. All data analyses and plots were generated using GraphPad Prism 9 version 9.4.0 (GraphPad Software Inc. CA, USA). Symbols represent p-values of different orders of magnitude, * $p<0.05$, † $p< 0.01$, ‡ $p<0.001$, § $p<0.0001$ and ¶ $p<0.00001$ vs. Control; and # $p<0.05$, | |$p<0.01$ vs. HFpEF-Placebo.

*Supporting Information (SI): Figures*

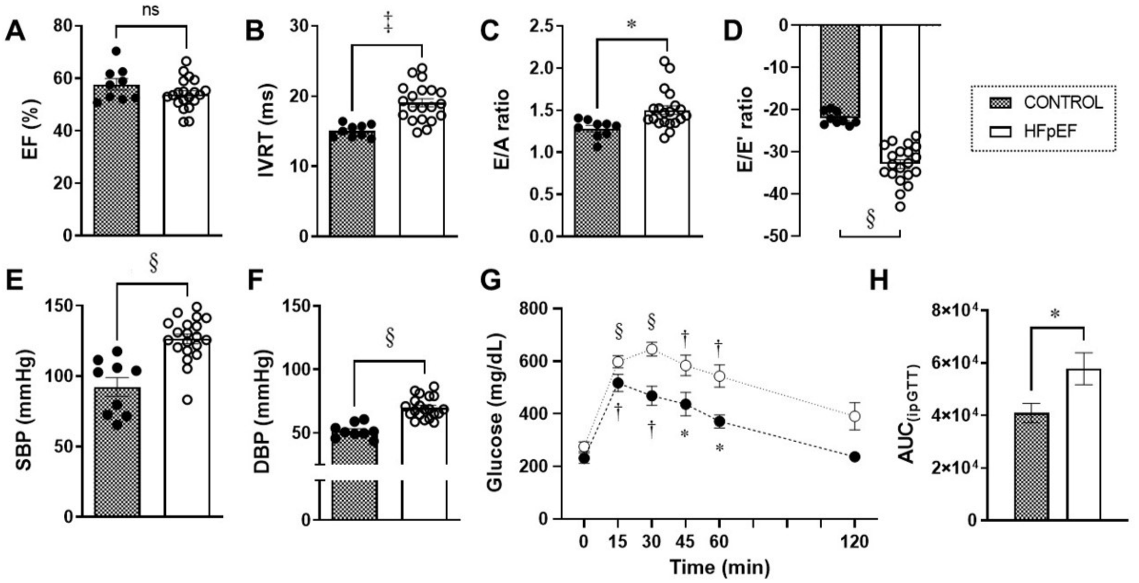


**Fig. S1. Baseline characteristics after 5 weeks of control and HFD plus L-NAME diet.** **(A)** There was no significant difference in ejection fraction (EF); however, **(B)** isovolumetric relaxation time (IVRT), **(C)** (E/A), and **(D)** (E/E') revealed impaired diastolic function in HFpEF mice compared to control mice. Both systolic **(E)** and diastolic **(F)** blood pressure were markedly increase in HFpEF mice compared to control (Unpaired t test: * $p<0.05$, ‡ $p<0.001$, § $p<0.0001$, n=9 for controls and n=20 for HFpEF). **(G)** Intraperitoneal glucose tolerance tests in control and HFpEF mice. (Two-way ANOVA for repeated measurements followed by Tukey's multiple comparisons' test * $p<0.05$, † $p<0.01$, § $p<0.0001$, n=10 for each group) and **(H)** area under the curve (AUC) of the ipGTT in HFpEF mice is increased in comparison to control suggesting glucose intolerance (Unpaired t test * $p<0.05$, n=10 for each group). All data are presented as mean ± SEM.

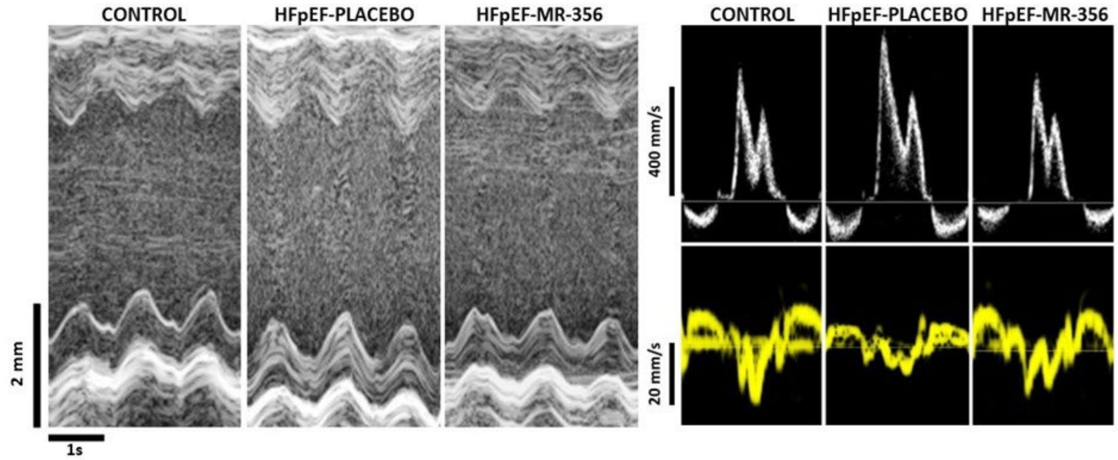


**Fig. S2. Echocardiographic images of control and HFpEF mice. (A)** Representative M-Mode tracings of parasternal short axis views, **(B)** mitral inflow pattern and **(C)** mitral annular velocity of control, HFpEF-placebo and HFpEF-MR-356 mice.

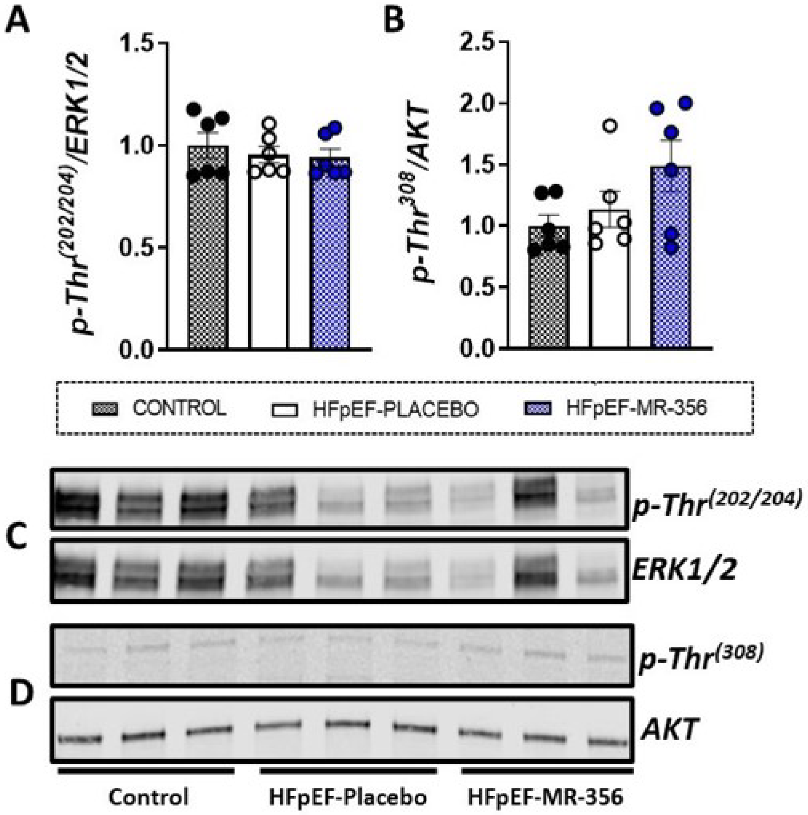


**Fig. S3.** Western blot analysis of **(A)** *p-Thr*$^{(202/204)}$ ERK1/2, and **(B)** *p-Thr* $^{(308)}$AKT expression normalized to the total ERK1/2, and AKT abundance in left ventricle homogenates did not show differences among the experimental groups (n=6). Representative immunoblots are shown in **C** and **D**, respectively. All data are presented as mean ± SEM.

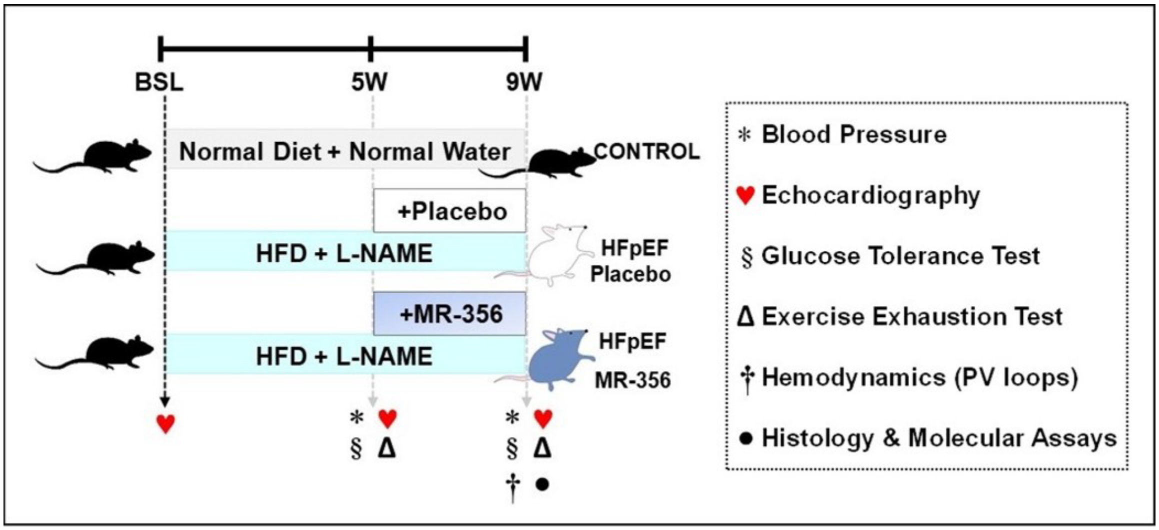


**Fig. S4. Experimental design.** C57BL6N (Charles River) male mice (8-weeks-old) were fed a high-fat diet (HFD, D12492, Research Diets, Inc) plus the nitric oxide synthase (NOS) inhibitor, $N^{\omega}$ – nitro-L-arginine methyl ester (L-NAME, 0.5 g/L, Sigma Aldrich #N5751) for 9 weeks to induce HFpEF (HFD+L-NAME). Food and water were replaced twice a week. After 5 weeks of HFD+L-NAME regimen, mice were randomly assigned to receive subcutaneous injections (100 µl) of either vehicle (DMSO + propylene glycol, HFpEF-placebo) or MR-356 (HFpEF-MR-356, 200 µg/kg) once a day for 4 weeks. Control animals received regular chow and water. Blood pressure, echocardiography, glucose tolerance and exercise exhaustion test were assessed at 5 and 9 weeks after beginning the HFD+L-NAME regimen. Hemodynamic evaluation and tissues collection for histology and molecular assays were obtained at the endpoint.

*Supporting Information* (*SI*)*: Tables*

**Table S1.** Daily averaged food intake and water intake from control mice (normal chow and no treatment) and HFpEF (HFD + L-NAME) mice treated with placebo or GHRH-MR-356 during a 9-week regimen.

| **Parameters** | **CONTROL (N=9)** | **HFpEF PLACEBO (N=10)** | **HFpEF MR-356 (N=10)** | ***P value*** |
|---|---|---|---|---|
| Food Intake (g/day) | 3.6±0.003 | 2.2±0.05 ¶ | 2.2±0.06 ¶ | ¶ p<0.00001 vs. Control |
| Water Intake (ml/day) | 3.8±0.03 | 2.5±0.05 § | 2.5±0.06 § | § p<0.0001 vs. Control |

All data are presented as mean ± SEM.

One-way ANOVA followed by Tukey's test: ¶ p<0.00001 vs. Control.

Kruskal-Wallis followed by Dunn's test: § p<0.0001 vs. Control.

**Table S2.** Animal's morphological characteristics post-mortem.

| Parameters | CONTROL (N=9) | HFpEF PLACEBO (N=10) | HFpEF MR-356 (N=10) | *P value* |
|---|---|---|---|---|
| **BW (g)** | 28.7±0.7 | 37.4±1.4 § | 35.2±1.2 † | † p<0.01 § p<0.0001 vs. Control |
| **TL (mm)** | 17.8±0.07 | 17.8±0.09 | 17.9±0.09 | ns |
| **HW (mg)** | 117.9±1.9 | 131.1±2.8 † | 127.5±3.1* | * p<0.05 † p<0.01 vs. Control |
| **LA (mg)** | 2.3±0.14 | 3.7±0.27 † | 2.7±0.15 # | # p<0.05 vs. Placebo † p<0.01 vs. Control |
| **LW (mg)** | 121.9±0.8 | 137.0±3.1 ‡ | 135.1±2.7 † | † p<0.01 ‡ p<0.001 vs. Control |
| **HW/TL (mg/mm)** | 6.6±0.1 | 7.4±0.1 † | $7.1±0.1^{0.055}$ | † p<0.01 vs. Control p=0.055 vs. Placebo |
| **LA/TL (mg/mm)** | 0.13±0.08 | 0.21±0.015 † | 0.15±0.08 \|\| | † p<0.01 vs. Control \|\| p<0.01 vs. Placebo |
| **LW/TL (mg/mm)** | 6.8±0.005 | 7.7±0.16 ‡ | 7.5±0.15 † | † p<0.01 ‡ p<0.001 vs. Control |
| **LWC (mg)** | 93.2±2.3 | 102.4±2.5 * | $101.4±2.2^{0.051}$ | * p<0.05 vs. Control p=0.051 vs. Placebo |

Body weight (BW), tibia length (TL), heart weight (HW), left atrium (LA) weight, lung weight (LW), heart weight to tibia length ratio (HW/TL), left atrium to tibia length (LA/TL), lung weight to tibia length ratio (LW/TL) and lung water content (LWC).

All data are presented as mean ± SEM.

One-Way ANOVA followed by Tukey's comparison test: * p<0.05, † p<0.01, ‡ p<0.001, § p<0.0001 vs. Control; and # p<0.05, || p<0.01, p=0.055, and p=0.051 vs. HFpEF-placebo.

**Table S3.** Blood pressure measurements from control mice and HFpEF mice treated with placebo or GHRH-MR-356.

| Parameters | CONTROL (N=9) | | HFpEF (N=20) | HFpEF PLACEBO (N=10) | HFpEF MR-356 (N=10) | *P value* |
|---|---|---|---|---|---|---|
| | 5 weeks | 9 weeks | 5 weeks | 9 weeks | 9 weeks | |
| **SBP (mmHg)** | 92±6.6 | 110±1.1 | 126±3.4§ | 126±1.9§ | 125±2.4§ | §p<0.001 vs. Control (at same time point) |
| **DBP (mmHg)** | 51±1.9 | 52±2.5 | 70±1.9§ | 67±2.9† | 70±2.5‡ | †p<0.01 ‡p<0.001 §p<0.0001 vs. Control (at same time point) |
| **MAP (mmHg)** | 71±1.7 | 71±2.1 | 90±1.8§ | 86±2.5‡ | 88±2.2§ | ‡p<0.001 §p<0.0001 vs. Control (at same time point) |

Systolic blood pressure (SBP), diastolic blood pressure (DBP), Mean arterial pressure (MAP).

All data are presented as mean ± SEM.

One-way ANOVA followed by Tukey's test: † $p<0.01$, ‡ $p<0.001$, § $p<0.0001$ vs. Control at the same time point.

**Table S4.** Changes in echocardiographic measurements at 5 and 9 weeks after HFD+L-NAME diet.

| Parameters | CONTROL (N=9) | | HFpEF (N=20) | HFpEF PLACEBO (N=10) | HFpEF MR-356 (N=10) |
|---|---|---|---|---|---|
| | **5 weeks** | **9 weeks** | **5 weeks** | **9 weeks** | **9 weeks** |
| **M-Mode** | | | | | |
| **HR (bpm)** | 513.5±11.7 | 540±15.4 | 528±9.6 | 515.9±10.1 | 568±20.2 |
| **LVESD (mm)** | 2.7±0.1 | 2.8±0.1 | 2.7±0.1 | 2.9±0.1 | 2.6±0.1 # |
| **LVEDD (mm)** | 3.8±0.01 | 3.9±0.1 | 3.7±0.1 | 4.1±0.1 | 3.7±0.1 # |
| **EF (%)** | 57.8±1.5 | 54.8±1.0 | 54.8±0.9 | 56.3±1.8 | 57.7±1.8 |
| **FS (%)** | 29.6±1.0 | 54.8±1.0 | 27.9±0.6 | 54.8±1.2 | 30±1.2 |
| **LV Mass (mg)** | 97.9±3.1 | 96.8±4.4 | 117.1±2.1 § | 146.7±6.2 § | 121.7±5.3 |
| **LV Mass Cor (mg)** | 78.3±2.5 | 77.5±3.5 | 93.7±1.7 § | 117.4±4.9 § | 97.3±4.3 |
| **LVAW;s (mm)** | 1.3±0.02 | 1.2±1.0 | 1.3±0.02 | 1.3±0.05 | 1.4±0.04 * |
| **LVAW;d (mm)** | 0.85±0.03 | 0.8±0.04 | 0.9±0.02 | 0.9±0.04 | 1.0±0.02 |
| **LVPW;s (mm)** | 0.98±0.03 | 0.9±0.3 | 1.1±0.02 † | 1.2±0.4 † | 1.2±0.8 * |
| **LVPW;d (mm)** | 0.6±0.001 | 0.6±0.02 | 0.8±0.02 § | 0.9±0.03 ‡ | 0.8±0.06 † |
| **RWTd** | 0.34±0.02 | 0.3±0.02 | 0.4±0.02 ‡ | 0.5±0.02 † | 0.4±0.03 * |
| **LVRI** | 20.6±0.5 | 19.8±.0.8 | 25.1±0.4 § | 28.8±0.92 § | 26.3±1.28 * |
| **B-mode** | | | | | |
| **HR (bpm)** | 540±9.1 | 564±12.1 | 555±7.5 | 548±12.5 | 566±16.4 |
| **ESV (µl)** | 17.7±1.1 | 21.9±0.7 | 18.5±0.7 | 20.5±1.3 | 19.2±0.9 |
| **EDV (µl)** | 43.9±1.5 | 48.3±1.5 | 42.6±1.3 | 46.7±2.0 | 43.1±1.7 |
| **SV (µl)** | 26.3±0.9 | 26.4±0.9 | 24.1±0.8 | 26.2±1.0 | 23.9±1.0 |
| **EF (%)** | 57.7±2.1 | 54.6±0.7 | 53.9±1.3 | 56.4±1.4 | 55.4±0.9 |
| **CO (µl)** | 13.4±0.5 | 14.9±0.6 | 13.0±0.3 | 14.5±0.8 | 13.7±0.6 |
| **Mitral Doppler** | | | | | |
| **E' (mm/s)** | -26.3±0.8 | -15.4±5.3 | -19.1±0.5 § | -9.3±4.5 † | -11.4±4.9 |

| | | | | | |
|---|---|---|---|---|---|
| **IVRT (ms)** | 15.0±0.3 | 14.2±0.6 | 19.3±0.5 ‡ | 19.3±0.8 ‡ | 18.27±0.8 † |
| **A (mm/s)** | 461±23.6 | 392±15.2 | 437±16.3 | 421±28.4 | 405±14.6 |
| **E (mm/s)** | 588.5±16.4 | 559±15.9 | 633±21.4 | 609±23.9 | 561±13.4 |
| **MPI** | 0.6±0.002 | 0.6±0.02 | 0.7±0.02 ‡ | 0.7±0.03 † | 0.7±0.04 † |
| **E/A** | 1.3±0.04 | 1.4±0.05 | 1.5±0.1 * | 1.5±0.1 | 1.4±0.1 |
| **E/E'** | –22.5±0.7 | –22.0±0.6 | –32.2±0.8 § | –34.3±1.3 ‡ | –27.8±1.3 † ‖ |

Heart rate (HR), end-diastolic volume (EDV), end-systolic volume(ESV), ejection fraction (EF), stroke volume (SV), LV end-diastolic diameter (LVEDD), LV end-systolic diameter (LVESD), relative wall thickness in diastole (RWTd), LV remodeling index (LVRI), left ventricle mass (LV mass), peak early diastolic mitral annular velocity (E'), isovolumetric relaxation time (IVRT), peak velocity of late filling (A wave), peak velocity of early filling (E wave), myocardial performance index (MPI), ratio of peak velocity of early to late filling of mitral inflow (E/A), ratio between E wave and peak early diastolic mitral annular velocity (E/E').

All data are presented as mean ± SEM.

One-way ANOVA followed by Tukey's multiple comparisons test or Kruskal-Wallis followed by Dunn's multiple comparisons test: * p<0.05, † p<0.01, ‡ p<0.001, § p<0.0001 vs. Control at the same time point, # p<0.05, ‖ p<0.01 vs. HFpEF-placebo at the same time point).

**Table S5.** Hemodynamic parameters

| Parameters | CONTROL N=9 | HFpEF PLACEBO (N=9) | HFpEF MR-356 (N=8) | *P value* |
|---|---|---|---|---|
| **Heart rate (bpm)** | 523 ± 8 | 513 ± 11 | 513 ± 12 | ns |
| | | ***Integrated Performance*** | | |
| **EF (%)** | 57 ± 1.8 | 60.2 ± 2.2 | 59.2 ± 3.2 | ns |
| **SW (mmHg x ml)** | 1928 ± 143 | 2285 ± 84.6 | 2260 ± 194.2 | ns |
| **SV(ml)** | 26.7 ± 1.5 | 26.9 ± 0.9 | 20.7 ± 1.8 | ns |
| **CO (ml/min)** | 13.9 ± 0.7 | 13.8 ± 0.6 | 12.9 ± 0.8 | ns |
| **Ea/Ees** | 1.6 ± 0.2 | 1.4 ± 0.3 | 1.6 ± 0.4 | ns |
| | | ***Afterload*** | | |
| **ESP (mmHg)** | 84.1 ± 3.0 | 102.5 ± 3.7 ‡ | 97.2 ± 2.3 * | * p<0.05 ‡ p<0.001 vs. Control |
| **Ea (mmHg/ml)** | 3.2 ± 0.2 | 3.9 ± 0.2 | 4.0 ± 0.4 | ns |
| | | ***Preload*** | | |
| **EDP (mmHg)** | 5.7 ± 0.4 | 10.2 ± 1.7 * | 4.9 ± 0.5 # | *p<0.05 vs. Control #p<0.05 vs. Placebo |
| **EDV(ml)** | 46.6 ± 1.7 | 45.2 ± 1.6 | 42 ± 2.5 | ns |
| | | ***Contractility*** | | |
| **$dP/dT_{max}$ (mmHg/s)** | 7241 ± 300 | 9309 ± 606 † | 9879 ± 376 † | †p<0.01 vs. Control |
| **$dP/dT_{max}$_EDV (mmHg/s per ml)** | 140.8 ± 19.6 | 155.7 ± 21.0 | 178.3 ± 27.2 | ns |
| **Ees (mmHg/ml)** | 2.1 ± 0.2 | 3.4 ± 0.5 | 3.0 ± 0.4 | ns |
| **PRSW (mmHg)** | 52.7± 5.7 | 67.2 ± 12.0 | 58.4 ± 5.7 | ns |
| | | ***Lusitropy*** | | |
| **$dP/dT_{min}$ (mmHg/s)** | -6163 ± 396 | -8486 ± 564 * | -8793 ± 284 † | * p<0.05 † p<0.01 vs. Control |
| **EDPVR (exponential)** | 0.02 ± 0.003 | 0.03 ± 0.004 † | 0.02± 0.002 # | †p<0.01 vs. Control #p<0.05 vs. Placebo |
| **Tau (ms)** | 6.5 ± 0.3 | 7.0 ± 0.005 | 5.8 ± 0.2 | ns |

Heart rate (HR), ejection fraction (EF), stroke work (SW), stroke volume (SV), cardiac output (CO), arterial elastance (Ea) and slope of end-systolic pressure volume relationship (Ees) ratio (Ea/Ees), left ventricle end-systolic pressure (ESP), arterial elastance (Ea), end-diastolic pressure (EDP), left ventricle end-diastolic volume (EDV), maximal rate of pressure rise (dP/dTmax), relation of maximal rate of pressure rise and end diastolic volume (dP/dTmax_EDV), end-systolic pressure volume relationship [ESPVR](Ees), preload recruitable stroke work (PRSW), maximal rate of pressure decline (dP/dTmin), end-diastolic pressure volume relationship (EDPVR), relaxation time constant calculated by Weiss method (Tau).

All data are presented as mean ± SEM.

One-way ANOVA followed by Tukey's multiple comparisons test or Kruskal-Wallis followed by Dunn's multiple comparisons test, * $p<0.05$, † $p<0.01$, ‡ $p<0.001$vs. Control, # $p<0.05$ vs. HFpEF-placebo.

**Table S6.** Antibodies list

| Western Blotting | | | | | |
|---|---|---|---|---|---|
| **Primary Antibodies** | **Company** | **Catalog #** | **Description** | **Dilution** | **MW (kDa)** |
| Akt | CST | 9272 | Rabbit | 1/1000 | 60 |
| phospho Akt (Thr308) | CST | 2965 | Rabbit | 1/1000 | 60 |
| ERK 1/2 | CTS | 4695 | Rabbit | 1/1000 | 42-44 |
| phospho ERK 1/2 | CST | 9106 | Mouse | 1/2000 | 42-44 |
| iNOS | CST | 1312 | Rabbit | 1/1000 | 131 |
| IRE-1$\alpha$ | CST | 3294 | Rabbit | 1/1000 | 130 |
| phospho IRE-1a (Ser724) | Novus Biological | NB100-2323 | Rabbit | 1/1000 | 130 |
| MyBPC3 | Santa Cruz | sc-137237 | Mouse | 1/1000 | 144 |
| phospho MyBPC3 (Ser282) | Enzo | ALX-215-057-R050 | Rabbit | 1/5000 | 150 |
| Pro-BNP | Abcam | Ab239514 | Mouse | 1/500 | 22 |
| Troponin I | Santa Cruz | sc-365446 | Rabbit | 1/1000 | 37 |
| phospho Troponin I (Ser 23/24) | CST | 4004 | Rabbit | 1/1000 | 28 |
| phospho Troponin I (Ser43) | PhosphoSolutions | p2010-43 | Rabbit | 1/2000 | 25 |
| VEGF-A | Abcam | ab46154 | Rabbit | 1/1000 | 25 |
| GAPDH | CST | 97166 | Mouse | 1/10000 | 37 |
| GAPDH | CST | 2118 | Rabbit | 1/10000 | 4 |
| **Secondary Antibodies** | | | | | |
| IRDye 680RD | LiCor | 925-68071 | Rabbit | 1/15000 | |
| IRDye 800CW | LiCor | 925-32210 | Mouse | 1/15000 | |
| **Immunofluorescence** | | | | | |
| Isolectin B4-Alexa Fluor ™ 468Conjugate | Thermo Fischer | GS-IB4 (I21411) | Griffonia Simplicifolia | 1/50 | |
| DAPI (4′,6-diamidino-2-phenylindole) | Sigma-Aldrich | D9542 | Nucleic acid staining | 1/2000 | |

***Supporting Information (SI): References***


1. C. National Research Council Committee for the Update of the Guide for the, A. Use of Laboratory, "The National Academies Collection: Reports funded by National Institutes of Health" in Guide for the Care and Use of Laboratory Animals. (National Academies Press (US) Copyright © 2011, National Academy of Sciences., Washington (DC), 2011), 10.17226/12910.
